# Local microtubule and F-actin distributions fully determine the spatial geometry of *Drosophila* sensory dendritic arbors

**DOI:** 10.1101/2023.02.24.529978

**Authors:** Sumit Nanda, Shatabdi Bhattacharjee, Daniel N. Cox, Giorgio A. Ascoli

## Abstract

Dendritic morphology underlies the source and processing of neuronal signal inputs. Morphology can be broadly described by two types of geometric characteristics. The first is dendrogram topology, defined by the length and frequency of the arbor branches; the second is spatial embedding, mainly determined by branch angles and tortuosity. We have previously demonstrated that microtubules and actin filaments are associated with arbor elongation and branching, fully constraining dendrogram topology. Here we relate the local distribution of these two primary cytoskeletal components with dendritic spatial embedding. We first reconstruct and analyze 167 sensory neurons from the *Drosophila* larva encompassing multiple cell classes and genotypes. We observe that branches with higher microtubule concentration are overall straighter and tend to deviate less from the direction of their parent branch. F-actin displays a similar effect on the angular deviation from the parent branch direction, but its influence on branch tortuosity varies by class and genotype. We then create a computational model of dendritic morphology purely constrained by the cytoskeletal composition imaged from real neurons. The model quantitatively captures both spatial embedding and dendrogram topology across all tested neuron groups. These results suggest a common developmental mechanism regulating diverse morphologies, where the local cytoskeletal distribution can fully specify the overall emergent geometry of dendritic arbors.

## Introduction

Nervous systems comprise a set of distinct neuron types with specific functional roles. During development, each neuron matures into a type-specific axonal-dendritic architecture through a combination of genetic encoding and external influences, such as molecular guidance cues. The overall geometry of the dendritic arbor forms the basis of the neuron’s connectivity and computational properties (Jan and Jan, 2010; Lefebvre, 2021; Lefebvre et al., 2015; Parekh and Ascoli, 2015). The extensive influence of dendritic structure over neural function is further revealed by the experimentally observed correlations between altered neural morphology and various neurological disorders (Emoto, 2011; Falke et al., 2003; Forrest et al., 2018).

The morphology of dendritic arbors can be conceptualized in terms of two distinct geometric aspects: dendrogram topology and spatial embedding (Brown et al., 2008). Dendrogram topology defines tree size (total length) and complexity (number of branches), and affects the biophysical computations carried out by dendrites (Komendantov and Ascoli, 2009). Spatial embedding determines the overall shape of dendritic arbors and influences their connectivity to either sensory or axonal inputs (Ropireddy and Ascoli, 2011; Stiso and Bassett, 2018). Both topological and spatial aspects are exquisitely regulated to optimize each neuron type-specific functional role.

Mature neuronal arbors emerge from a series of causal events such as growth, retraction, and branching (Castro et al., 2020; Nanda et al., 2018b). Since each event occurs in three-dimensional space, the orientations of the newly formed branches are also determined simultaneously. Hence, elucidating the biochemical mechanisms underlying these processes entails relating spatially oriented topological events with the mediator molecules that drive arbor growth and plasticity.

All regulatory processes that control dendritic development, such as cell-specific genetic information, extracellular cues, spatial constraints, and neural activity (Jan and Jan, 2010; Lefebvre et al., 2015), eventually converge on the cytoskeletal mediator molecules, namely actin filaments (F-actin) and microtubules (Bhattacharjee et al., 2022; Das et al., 2017; Franker and Hoogenraad, 2013; Ledda and Paratcha, 2017; Nagel et al., 2012; Nanda et al., 2020). Additionally, the modification and maintenance of type-specific neuronal morphology is greatly facilitated by the dynamic self-organization of microtubule and F-actin (Coles and Bradke, 2015; Kapitein and Hoogenraad, 2015). Cell-intrinsic signaling and combinatorial interactions of transcription factors mold type-specific dendritic diversity by regulating cytoskeletal composition (Corty et al., 2016; Das et al., 2021; de la Torre-Ubieta L. and Bonni, 2011; Iyer et al., 2012)

While numerous upstream signaling pathways that regulate both microtubules and F-actin dynamics have been described (Bhattacharjee et al., 2022; Das et al., 2021; Hely et al., 2001; Nanda et al., 2017), their specific roles in the causal steps that produce dendritic morphology are only partially understood. Previously we have demonstrated that, in sensory neurons from the *Drosophila* larva, microtubule concentration is correlated with downstream arbor length, while actin filaments are enriched at branch points. We also reported that an arbor generation model can accurately reproduce dendrogram topology by controlling key events such as elongation, branching, and termination solely based on local cytoskeletal composition (Nanda et al., 2020).

Here we reconstructed mature Class IV and Class I dendritic arborization (da) neuron morphologies from confocal image stacks containing arbor-wide signals for microtubule and F-actin quantity (Bhattacharjee et al., 2022; Das et al., 2021; Nanda et al., 2018a). We then evaluate whether these two major cytoskeletal components also influence the dendritic arbor spatial embedding along with its dendrogram topology. We observe that higher quantity of local microtubule is associated with reduced angular deviations and straighter branches, and report additional significance effects of F-actin. Informed by these correlations, we proceed to test whether local cytoskeletal composition alone is sufficient to reproduce all emergent geometric features of dendritic morphology, including spatial embedding. Hence, we create a computational model where the microtubule and F-actin distributions determine not only elongation and bifurcations, but also the local orientation of all branches.

## Methods

Multi-signal morphological reconstructions were acquired using previously described methods (Nanda et al., 2021, 2018a), as briefly summarized below. The processed data have been deposited in Mendeley.com along with the analysis and modeling code for open-access release upon publication of this manuscript (preview: data.mendeley.com/datasets/cj69j8bpn8/draft?a=b9d99a91-9048-4974-b946-f99c397a79ec). The same Mendeley.com package also includes all analysis and simulations scripts which are released open source to facilitate reproducibility and further community development (Gleeson et al., 2017). The digital tracings and enhanced ESWC files of all 167 neurons have been submitted to NeuroMorpho.Org (Akram et al., 2018) as part of the Ascoli and Cox archives (Table 1).

**Table 1:** Abbreviation, neuron class, subtype, genotype, morphological characteristics, number of cells, and data sources information for the 17 neuron groups used in this study.

| Abbreviation | Neuron Class | Neuron Subclass | Mutation | Morphology Description | Number of Neurons | Source Publication |
| --- | --- | --- | --- | --- | --- | --- |
| C1-ddaE-WT | Class I | ddaE | Wild type | Simple Dendritic Arbors Secondary arbors tend to extend towards one direction | 19 | doi: 10.3389/fnmol.2022.926567 |
| C1-vpda-WT | Class I | vpda | Wild type | Simple Dendritic Arbors, Secondary arbors tend to extend towards both direction | 8 | doi: 10.3389/fnmol.2022.926567 |
| C1-vpda-mts-IR | Class I | vpda | mts- Knockdown | Reduction in Size and Complexity | 9 | doi: 10.3389/fnmol.2022.926567 |
| C1-vpda-mts-OE | Class I | vpda | mts-Overexpression | Slight reduction in size, complex terminal branches | 12 | doi: 10.3389/fnmol.2022.926567 |
| C4_WT | Class IV | ddaC | Wild type | Large Complex dendritic arbors | 10 | doi: 10.3389/fnmol.2022.926567, doi:10.1242/dev.187609 |
| C4_bdwf-IR | Class IV | ddaC | CG3995/bdwf Knockdown | Reduced span, small terminal branches | 8 | doi: 10.1101/2023.02.15.528686 |
| C4_form3-OE | Class IV | ddaC | formin 3 Overexpression | Reduced size and complexity | 9 | doi:10.1242/dev.187609 |
| C4_mts-IR | Class IV | ddaC | mts- Knockdown | Reduction in size, complexity shifted to distal branches | 11 | 10.3389/fnmol.2022.926567 |
| C4_mts-OE | Class IV | ddaC | mts Overexpression | Reduced size and span | 10 | 10.3389/fnmol.2022.926567 |
| C4_cut_IR | Class IV | ddaC | cut Knockdown | Reduced size, reduction in terminal complexity | 8 | doi: 10.1101/2023.02.15.528686 |
| C4_RpL4-IR | Class IV | ddaC | RpL4- Knockdown | Reduced size and major reduction in complexity | 10 | doi: 10.1101/2023.02.15.528686 |
| C4_RpL17-IR | Class IV | ddaC | RpL17- Knockdown | Reduced size and complexity | 6 | doi: 10.1101/2023.02.15.528686 |
| C4_RpL22-IR | Class IV | ddaC | RpL22- Knockdown | Reduced complexity | 10 | doi: 10.1101/2023.02.15.528686 |
| C4_RpL31-IR | Class IV | ddaC | RpL31- Knockdown | Reduced size and complexity | 9 | doi: 10.1101/2023.02.15.528686 |
| C4_RpL35A-IR | Class IV | ddaC | RpL35A- Knockdown | Reduced size | 13 | doi: 10.1101/2023.02.15.528686 |
| C4_RpS10b-IR | Class IV | ddaC | RpS10b- Knockdown | Reduced size | 8 | doi: 10.1101/2023.02.15.528686 |
| C4_RpS24-IR | Class IV | ddaC | RpS24- Knockdown | Reduced size | 7 | doi: 10.1101/2023.02.15.528686 |

### *Drosophila* strains and live confocal imaging

*Drosophila* stocks were reared at 25°C on standard cornmeal-molasses-agar media. Age-matched wandering third instar larvae of both sexes were used for all experiments. Fly strains used in this included: *UAS-GMA::GFP; GAL4[477],UAS-mCherry::Jupiter* (Das et al., 2017) (Class IV cytoskeletal reporter strain); *UAS-GMA::GFP;+;GAL4[221],UAS-mCherry::Jupiter (Class* I cytoskeletal reporter strain) (Bhattacharjee et al., 2022); *UAS-form3-B1* (Tanaka et al., 2004), *UAS-bdwf-IR (B27997), UAS-mts-IR (B57034), UAS-mts (B53709), UAS-ct-IR (B33965), UAS-RpL4-IR (v101346), UAS-RpL17-IR (v105376), UAS-RpL22-IR (v104506), UAS-RpL31-IR (v104467), UAS-RpL35A-IR (v100797), UAS-RpS10b-IR (v106323), UAS-RpS24-IR (v104676). Oregon R* was used as a genetic background control for outcrosses to the multi-fluorescent cytoskeletal transgene reports. We previously confirmed that expression of the F-actin and microtubule transgene reporters did not themselves exert any effects on dendritic development (Das et al., 2017). Virgin flies from either *UAS-GMA; GAL4[221], UAS-mCherry::JUPITER,* or *UAS-GMA; GAL4[477], UAS-mCherry::Jupiter were* crossed to gene-specific *UAS-RNAi* transgenic males. UP-TORR (www.flyrnai.org/up-torr/) was utilized to evaluate predicted off-target effects for gene-specific RNAi transgenes (Hu et al., 2013) all RNAi lines used in this study have no predicted off-target effects. Live confocal imaging of age-matched third instar larval mature da neuron Class I and Class IV arbors was performed as previously described (Das et al., 2017; Nanda et al., 2018b, 2018a) Briefly, larvae were placed on a glass slide and immersed in a 1:5 (v/v) diethyl ether to halocarbon oil and covered with a 22×50 mm cover slip. Neurons expressing fluorescent protein cytoskeletal reporter transgenes were visualized on a Zeiss LSM 780 confocal microscope. Images were collected as z-stacks at a step size of 1-2 microns and 1024 x 1024 resolution using a 20X air objective (Plan Apo M27, NA 0.8) and 1 airy unit for the pinhole.

### Morphological reconstruction and editing

Digital reconstructions of neuronal morphology are lists of interconnected compartments, each representing a small section of neurite between two consecutive tracing locations (Nanda et al., 2018a). Dendritic morphology was semi-automatically reconstructed and edited from the two-channel confocal images using neuTube (Feng et al., 2015). A third channel (the pseudo membrane), created by combining microtubule and F-actin signals in FIJI (Schindelin et al., 2012), was used for tracing the primary reconstruction. Next, any inaccuracies in tree topology were resolved with the TREES Toolbox (Cuntz et al., 2010). The resultant files were then manually curated in neuTube once more.

Each neuron was used as input in the Vaa3D ***Multichannel_Compute*** plug-in tool (run in Vaa3D version 3.1) to capture signal information from both the microtubule (MT) and F-actin substructures into the final multi-signal reconstructions. The local cytoskeletal quantity (CQ) of MT and F-actin signal within a compartment was quantified using the following equation.

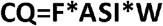

Here, F is the fraction of the dendritic compartment volume occupied by the signal, ASI is the average signal intensity, and W is local dendrite thickness, which is approximated by measuring the local diameter of the compartment. All coordinates were then multiplied by the voxel size in physical units (μm) to generate the final multi-signal reconstructions. Microtubule or F-actin quantities are referred to as just microtubule or F-actin in the remaining text for simplicity.

The integral microtubule (IM) of each compartment measures the microtubule quantity throughout the upstream arbor path, starting from the soma up to that compartment. IM for the (i+1)^th^ compartment is defined as:

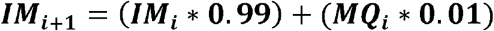

where MQ is the quantity of microtubule, and the i^th^ compartment is the parent of the (i+1)^th^ compartment.

All multi-signal reconstructions were programmatically processed to resolve any trifurcations and overlapping points. Additionally, all the dendritic branches were locally resampled so as to make each compartment 2 μm long.

### Measurement and correlation of morphological and cytoskeletal properties

Analyses were conducted both at the compartment level and branch level. Here, a dendritic branch is defined as a series of consecutive compartments beginning at a tree stem or bifurcation point and ending at the next bifurcation or at a termination point. MATLAB scripts were written to quantify, for every neuron, each compartment length, orientation, angular deviation, MT and F-actin quantities, path distance from the soma, branch order, and downstream arbor lengths. More than half a million dendritic compartments from all 17 neuron groups were measured for all the parameters. Corresponding branch level parameters were computed from the values of the constituent compartments. For example, average branch microtubule was calculated as the mean MT quantity of the compartments forming the branch. Branch straightness was defined as ratio of branch Euclidean length (i.e., straight line distance between branch start and end point) to branch path length (i.e., sum of length of all compartments in the branch). The remote branch angle was measured as the angle between a branch straight line relative to the straight line of its parent branch.

The correlation amongst the branch level parameters were calculated for each neuron group. If two variables were significantly correlated, the correlation coefficient of their binned averages was also computed. In these cases, the independent cytoskeletal parameter (e.g., microtubule quantity or F-actin quantity) values were placed into 20 bins of approximately equal sample size. The average of the dependent properties (e.g., branch remote angle or branch straightness) at each bin were then correlated (and plotted) against the bin averages of the independent cytoskeletal properties.

For each pair of ‘sibling’ branches originating from the same bifurcation point, we also measured the magnitude and direction of their deviations from the parent branch, with positive and negative signs assigned to counterclockwise and clockwise deviations, respectively. We define branch tilt as the angle created between the parent branch and the straight line that splits the two sibling branches through the middle, creating two angles of the same magnitude.

### Model of dendritic arbor generation

The full arbor generation model is built at the single compartment resolution level and extends our previous simulation of dendrogram topology (Nanda et al., 2020). At each iteration, the model reads the compartmental composition of microtubule and F-actin as input, and outputs the developmental event (extend, bifurcate or terminate) as well as the cytoskeletal composition and the angular deviation of the newly created child compartment(s), if any. As in our earlier model (Nanda et al., 2020), the process relies on three two-dimensional sampling grids, where the dimensions are local microtubule and F-actin quantities: event grid, extension grid, and branching grid. The event grid determines the probability of extending, bifurcating or terminating based on the compartment MT and F-actin composition. Similarly, the extension and branching grids determine the cytoskeletal composition and angular deviation of the newly created child compartment(s) also based on the (parent) compartment microtubule and F-actin quantities. Within each sampling grid bin, the model choses the topological event, cytoskeletal composition, and angular orientation that best match the integral microtubule of the parent compartment.

### Comparison of real and simulated arbor properties

Once virtual neuron arbors are generated, they are statistically compared against their real counterparts. First, the L-Measure (Scorcioni et al., 2008) morphometric tool is used to quantify 10 distinct properties for each real and simulated neuron. These 10 measures are then separately compared between real and simulated neuron groups using the student’s t-test (if N>10 neurons per group) or the non-parametric Wilcoxon Rank Sum test (if N<=10 per group).

Persistent vector analysis was carried out using a publicly available automated tool (Bijari et al., 2021), which automatically produce one persistent vector file for each neuronal reconstruction file. A new analysis script measured the arccosine distances between the persistence vectors of each neuron pair to compare within-group differences against between-group differences.

Dendritic density analysis was carried out using TREES Toolbox (Cuntz et al., 2010). Two-dimensional density matrices were created since the sensory da neurons lie flat on the cuticle of the *Drosophila* larva. These matrices were then vectorized and analyzed in terms of arccosine distances as described above for the persistence vectors.

## Results

We studied 167 *Drosophila melanogaster* larval da neurons from 17 distinct groups, including wild type (WT) controls and mutants in the simplest Class I and the most complex Class IV neurons. Four Class I groups included two WT subtypes (ddaE and vpda) and two mutants of the vpda subtype *(mts* knockdown and *mts* overexpression). Thirteen Class IV groups, all of subtype ddaC, included WT and 12 mutants: *bdwf* knockdown, *form3* overexpression, *mts* knockdown, *mts* overexpression, *ct* knockdown, *RpL4* knockdown, *RpL17* knockdown, *RpL22* knockdown, *RpL31* knockdown, *RpL35A* knockdown, *RpS10b* knockdown, and *RpS24* knockdown (Table 1).

### Distinct dendritic architecture of sensory neurons

The morphological appearance differed substantially between Class I and Class IV neurons as well as among genotypes. These differences in overall morphology were accompanied by changed microtubule and F-actin distributions across the dendritic arbors (Figure 1). Class I ddaE and Class I vpda subtypes also have distinctly different shapes. The ddaE neurons have comb-like secondary branches extending towards the same direction from the primary branch whereas the vpda secondary branches extend towards both sides. All Class IV ddaC mutants invade a reduced spatial surrounding compared to the WT control ddaC neuron, hinting that among the genetic perturbations we analyzed (irrespective of whether knockout or overexpression) the phenotypic defect leads to a less optimized space-filling architecture (Fig.1).

**Figure 1:**
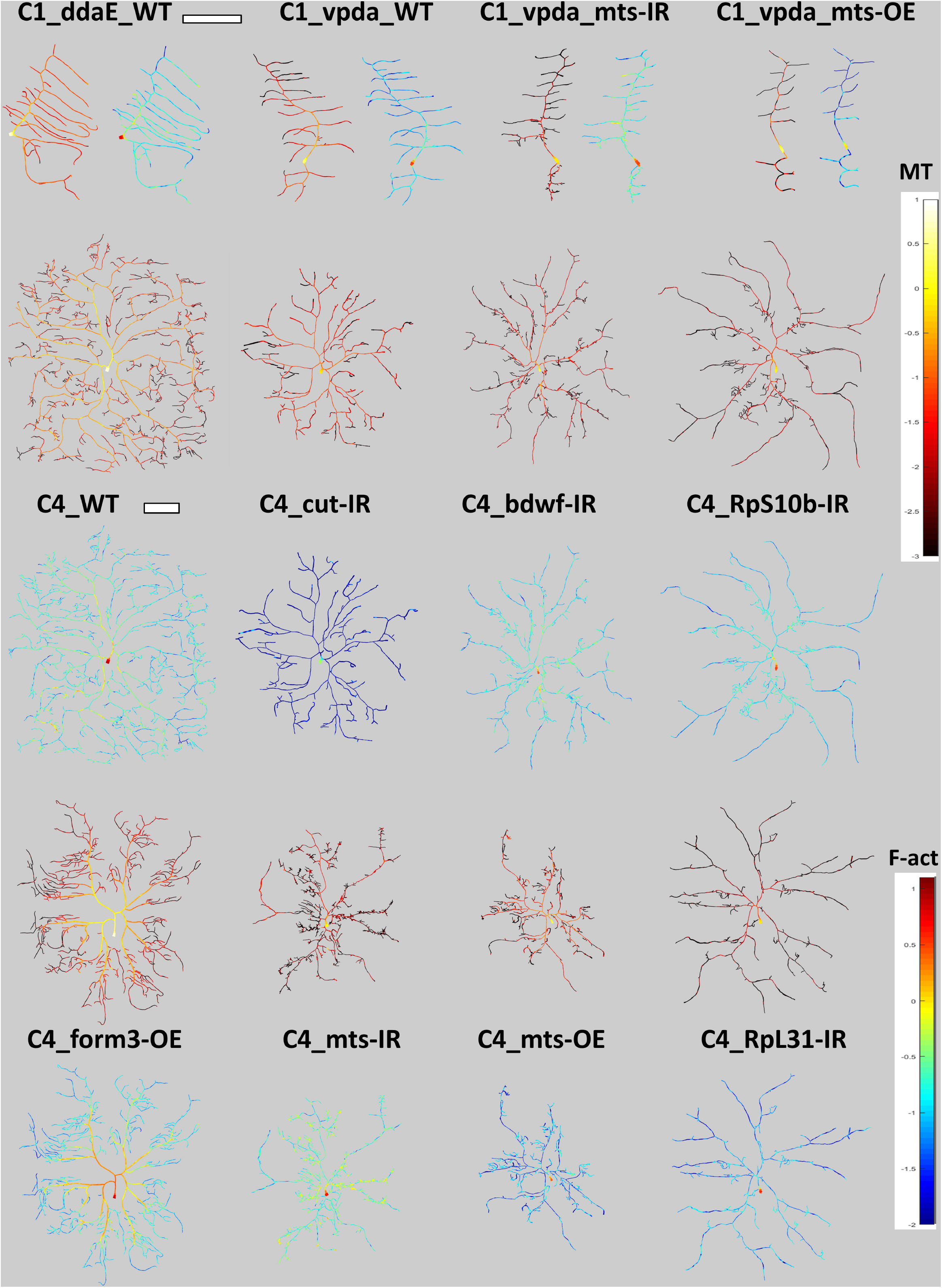
Morphological Diversity of Sensory Neurons in the *Drosophila* Larva. Single representative neurons from 4 Class I neuron types (1 WT subclasses and 2 mutant groups) and 8 Class IV neuron types (1 WT ddaC and 7 mutant groups) are shown, each in two separate images displaying microtubule (red hue) and F-actin (green-blue hue) distributions, respectively. Color bars are constant for all 17 neurons groups. Two separate scale bars for Class I and Class IV neuron groups each represent 100 microns.

We observe a distinctive morphological signature in the distributions of branch lengths and angles of *Drosophila* sensory neurons. Specifically, in *Drosophila* Class IV neurons, terminal branches are significantly (p < 0.05) shorter (12.9 ± 12.3 μm) than internal branches (17.6 ± 15.3 μm). Moreover, branch angles increase with branch order in both class I ddaE WT (binned correlation R = 0.17, p<0.05) and Class IV ddaC WT (binned correlation R = 0.05, p<0.05) neurons (see Suppl. Table 1 for corresponding data for all neuron groups). Interestingly, these architectural features are opposite to those reported in other neural systems (see Discussion).

### Influence of local cytoskeleton composition on branch orientation and tortuosity

In order to estimate the relative influence of cytoskeletal composition over spatial geometry, we measured the angular deviation (Fig. 2A) and straightness (Fig. 2B) of every branch in each neuron across all groups and computed their correlations with MT and F-actin quantities (see Methods for details). In all 17 neuron groups, the remote branch angle was strongly anti-correlated to the average branch MT quantity (Fig. 2C and Table 2). Moreover, 12 out of 17 neurons groups, including Class IV ddaC WT and Class I ddaE WT, demonstrated a positive correlation between branch straightness and average branch MT quantity (Fig. 2D and Table 2). This suggests that local microtubules not only influence the overall orientation of a dendritic branch relative to its parent branch, but also the extent of its tortuosity. For Class I vpda neuron groups (WT, *mts-IR,* and *mts-OE*), the only significant correlations were between average branch microtubule and remote branch angle (Fig. 2E). This could partly be explained by the fact that Class I vpda neuron branches are largely straight and lack much variability that could be explained by differential cytoskeletal distributions.

**Figure 2:**
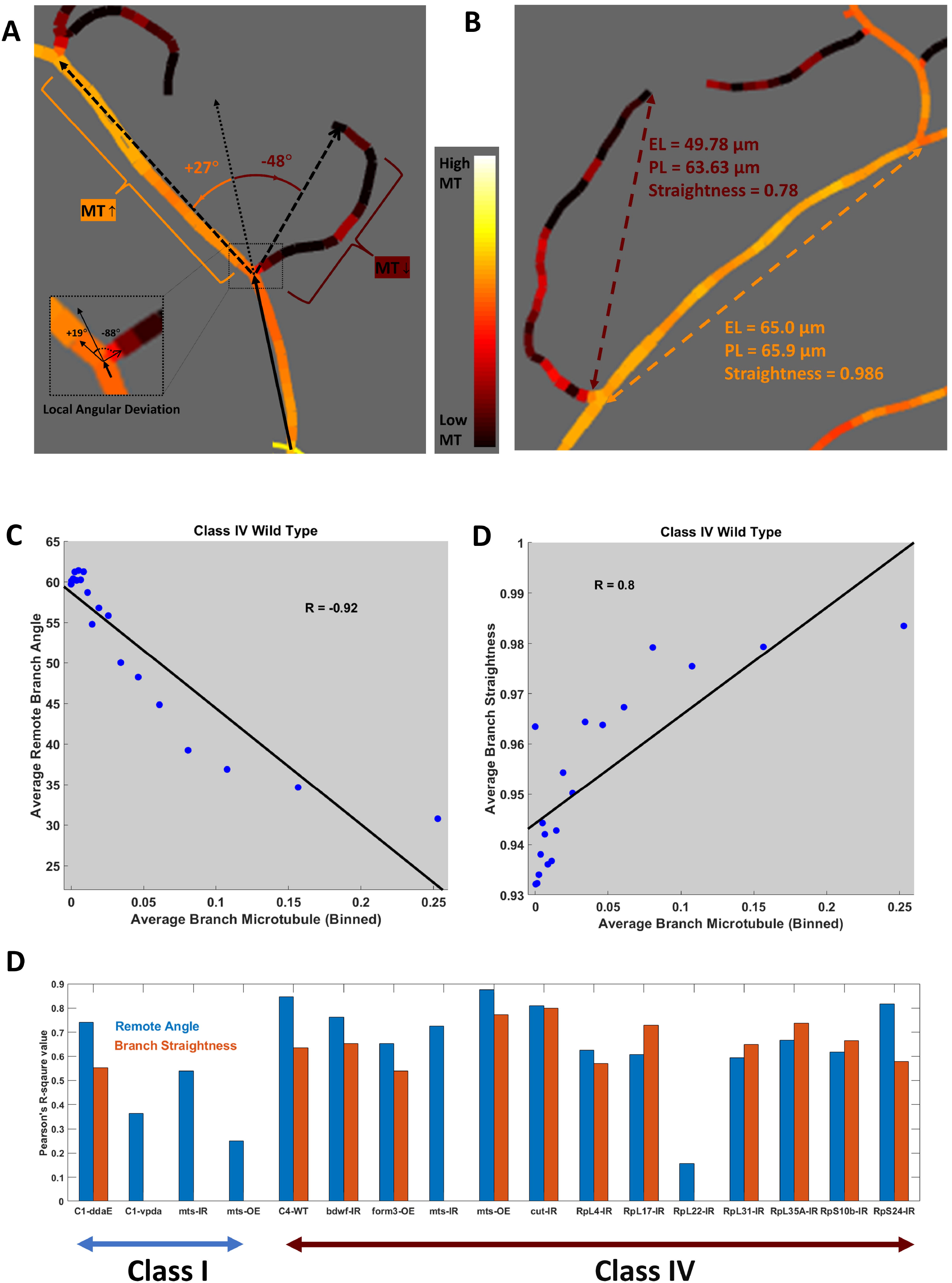
Branch Angular Deviation and Straightness. (A) Schematic illustration of remote angular deviation (inset: local angular deviation). (B) Schematic illustration of branch straightness (ratio of branch Euclidean length to branch path length). (C) Binned scatter plot of average remote branch angle against average branch microtubule quantity of Class IV wild type neurons. (D) Binned scatter plot of average branch straightness against average branch microtubule quantity of Class IV wild type neurons. (E) Coefficients of determination of branch MT with branch remote angle (blue) and straightness (purple). Class I vpda WT and mutants show no correlation between MT quantity and branch straightness. See Table 1 for sample size by genotype.

**Table 2:** Pearson’s correlations between average branch MT, F-actin or integral MT and remote branch angle or branch straightness for all 17 neuron groups. Values not reaching statistical significance after False Discovery Rate correction for multiple testing were substituted with 0s.

| Neuron Groups | MT vs. Remote Angle | F-act vs Remote Angle | Integral MT vs Remote Angle | MT vs. Branch Straightness | F-act vs. Branch Straightness | Integral MT vs. Branch Straightness |
| --- | --- | --- | --- | --- | --- | --- |
| C1-ddaE-WT | -0.79662 | -0.68174 | 0 | 0.64468 | 0 | 0.64137 |
| C1-vpda-WT | -0.40058 | 0 | 0 | 0 | 0 | 0 |
| C1-vpda-mts-IR | -0.57827 | 0 | 0 | 0 | 0 | 0 |
| C1-vpda-mts-OE | -0.23092 | 0 | 0 | 0 | 0 | 0 |
| C4_WT | -0.81964 | -0.88206 | -0.51383 | 0.72013 | 0.83697 | 0.10764 |
| C4_bdwf-IR | -0.73196 | 0 | 0 | 0.69391 | 0.68047 | 0 |
| C4_form3-OE | -0.71452 | -0.87823 | -0.6732 | 0.65887 | 0.85572 | 0.82909 |
| C4_mts-IR | -0.71502 | -0.63133 | 0 | 0 | 0.89429 | 0 |
| C4_mts-OE | -0.88264 | 0 | 0 | 0.82352 | 0.60876 | 0.45636 |
| C4_cut_IR | -0.74801 | 0.79775 | 0 | 0.84849 | 0 | 0.35218 |
| C4_RpL4-IR | -0.72041 | 0 | 0 | 0.63468 | 0.74358 | 0.53348 |
| C4_RpL17-IR | -0.61927 | 0 | -0.3319 | 0.71794 | 0.71844 | 0.32508 |
| C4_RpL22-IR | -0.14374 | 0 | -0.38076 | 0 | 0.53392 | 0 |
| C4_RpL31-IR | -0.6215 | 0 | 0 | 0.62481 | 0 | 0.49774 |
| C4_RpL35A-IR | -0.62188 | 0 | -0.53512 | 0.66933 | 0.4828 | 0.66447 |
| C4_RpS10b-IR | -0.59043 | -0.15194 | 0 | 0.67994 | 0.75233 | 0.49029 |
| C4_RpS24-IR | -0.79757 | -0.14115 | -0.45577 | 0.68023 | 0.54527 | 0.29519 |

Average branch F-actin quantity had a mixed correlation with remote branch angle across neuron types (Table 2). Similar to the average branch microtubule, in both Class IV ddaC WT and Class I ddaE WT neurons, as well as in four other groups (Class IV *form3-OE,* Class IV *mts-IR,* Class IV *RpS10b-IR,* and Class IV *RpS24-IR),* the correlation was negative (Table 2). In contrast, in Class IV *cut-IR* neuron subtype, average F-actin was positively correlated with remote angle, while the correlations were not statistically significant in the other groups. At the same time, average branch F-actin was positively correlated (like average microtubule) with branch straightness in 11 out of the 17 neurons groups, with the remaining ones, including all Class I groups, showing no significant correlation. Only in the case of Class IV *formin 3* overexpression, branch angle and straightness had stronger correlations with F-actin than with microtubule quantity.

Similar to local microtubule, integral microtubule was also negatively correlated with remote branch angle in 6 out of the 17 neuron groups (the remaining groups demonstrated no correlation) and was positively correlated with branch straightness in 11 out of 17 groups (Table 2).

### Relation between angular deviations of sibling branches

Local and remote branch angles were highly correlated across all neuron groups (Fig. 3 and Suppl. Table 2). This result demonstrates that angular orientations attained during branch formation tend to remain substantially preserved until the end of the branch (Fig. 3A,C). When we compare the sibling branch pairs, we also observe that the larger of the two sibling angles is highly correlated with the branch tilt (R = 0.88 for Class I ddaE WT, R = 0.94 for Class I vpda WT, R = 0.94 for Class IV ddaC WT). In most cases, moreover, the pairs have opposite signs of angular deviation. Additionally, the absolute angular deviation of the two sibling branches is negatively correlated (Fig. 3B,D): if the branch oriented counterclockwise relative to the parent branch deviates more from the parent direction, the branch oriented clockwise will deviate less, and vice versa (Suppl. Table 2). This angular homeostasis expands the types of morphological homeostasis previously observed both in *Drosophila* (Nanda et al., 2018b) and in the mammalian brain (Samsonovich and Ascoli, 2005).

**Figure 3:**
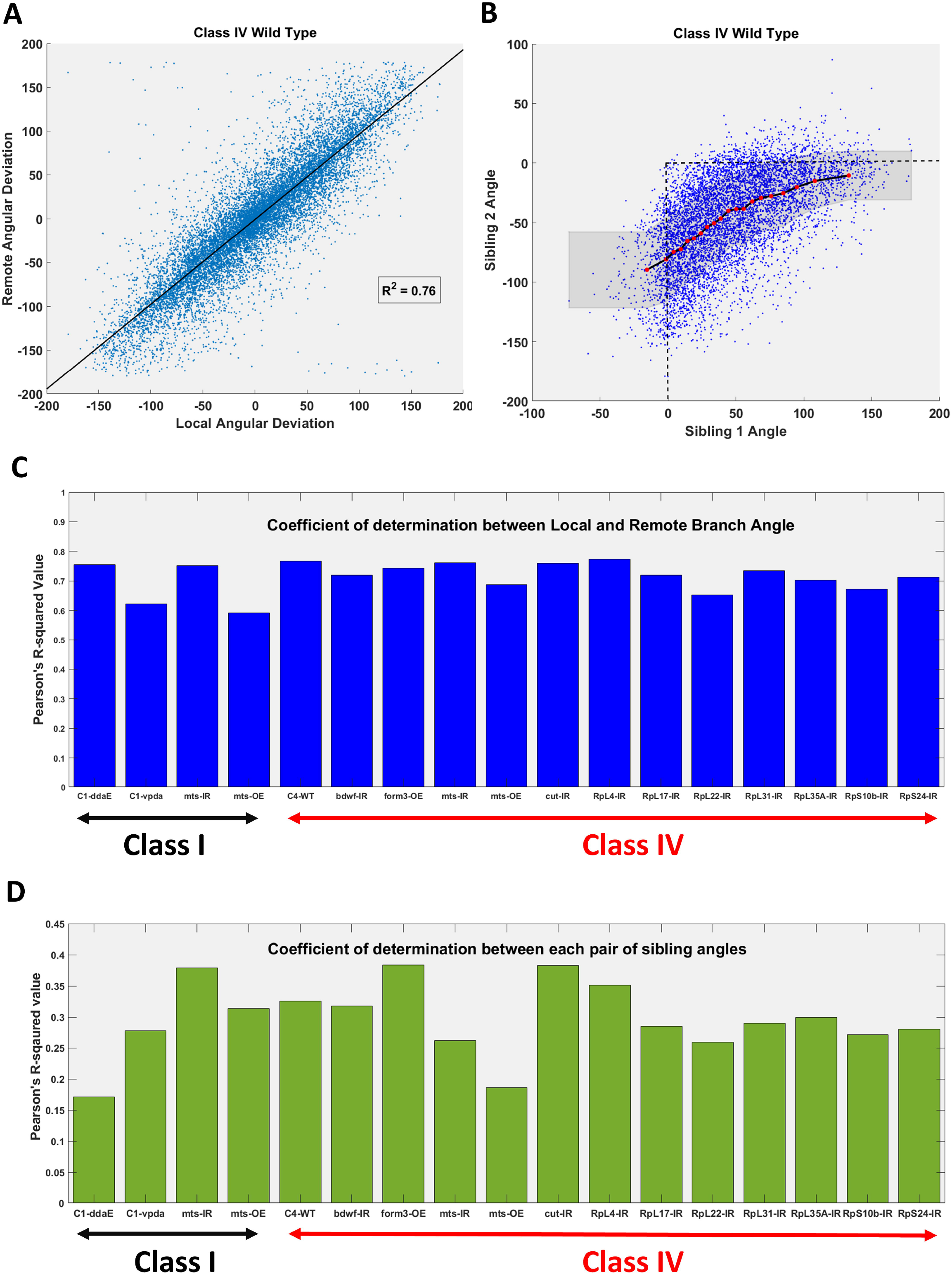
Analysis of Branch Angular Deviation. (A) Local and remote branch angles are strongly correlated in Class IV WT neurons. (B) The signed angular deviations of sibling branches are positively correlated when they deviate in opposite direction, corresponding to a negative correlation between the absolute angular deviations (see Suppl. Table 2) (C) Coefficients of determination between local and remote branch angles across the 17 neuron groups. (D) Coefficients of determination between each pair of sibling branches across the 17 neuron groups.

The above observations complement previous results implicating local MT and F-actin quantities in the quantitative specification of dendrogram topology (Nanda et al., 2020). Are the measured associations of cytoskeletal composition with branch angles and tortuosity, in addition to topology, sufficient to reproduce the complete arbor geometry of da neurons from various groups?

### A cytoskeleton-driven generative model of dendritic morphology

Every iterative step of the computational model for virtual tree generation consists of the simultaneous addition of one or two compartments at each growing branch (Fig. 4). Individual compartments provide the highest resolution of spatial embedding as well as cytoskeletal composition of a digital dendritic arbor. Even branch-level metrics such as straightness reflect an emergent property of the collective orientations of individual compartments. Thus, the compartment-level simulation not only allows for accurate local geometric embedding, but also more closely resembles the developmental process of dendritic trees. Informed by our experimental observations, the addition, cytoskeletal composition, and 3D embedding of newly generated compartments in the model are constrained by the local MT and F-actin quantities in the previous compartments, as sampled from real neurons.

**Figure 4:**
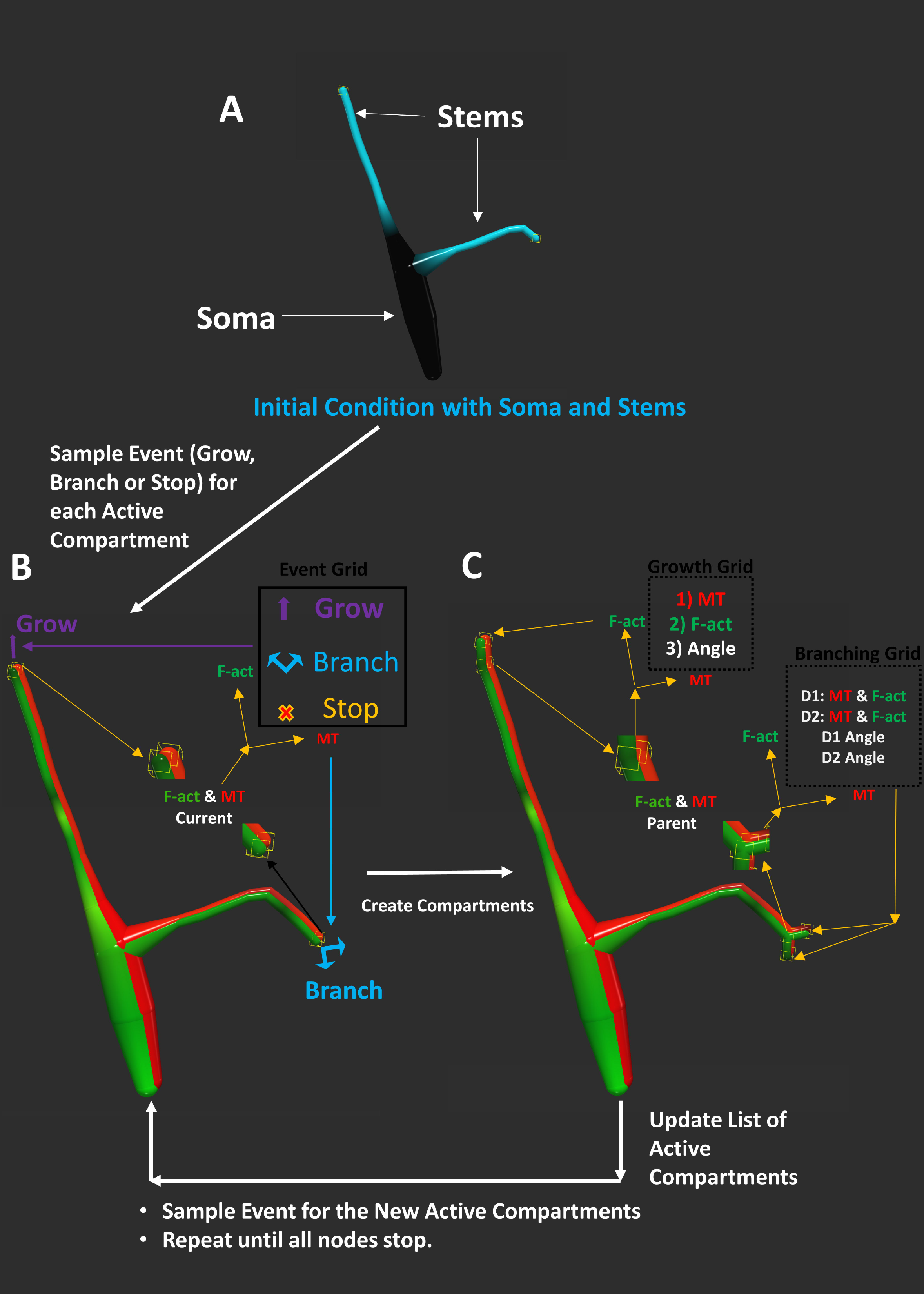
Computational Model of Dendritic Morphology. (A) The tree generation process starts with the initial MT and F-actin concentration of the soma and the dendritic stem end points as active compartments. (B) For each active compartment, the event sampling determines whether to grow (adding one new compartment), branch (adding two new compartments), or stop (ending the branch). (C) If a node grows or branches, the cytoskeletal composition of the newly created compartment is sampled from the corresponding data grids along with their angular deviations relative to their parent compartments. The process is repeated for all newly added compartments, until all branches stop.

The tree generation process starts with the MT and F-actin quantities of the soma and the dendritic stem end points as the active compartments (Fig. 4A). At each iteration, two sampling processes are carried out at every active compartment. The first sampling process (detailed in the Methods section) determines the topological fate (elongation, branching or termination) of the active compartment based on its MT and F-actin quantities (Fig. 4B), similar to our previous dendrogram topology model (Nanda et al., 2021). The second sampling process determines the cytoskeletal composition as well as the orientation of each newly generated compartment, also based on their parent compartment’s MT and F-actin composition (Fig. 4C). If a node elongates or bifurcates, the newly created compartments then become the new active compartments, and the sampling process iteration continues until all branches terminate. The same arbor generation model is employed for all 17 neurons groups.

### Simulated neurons reproduce all morphological attributes across neuron groups

To evaluate the effectiveness of the tree generation model, the simulated virtual neurons were first compared visually against their real counterparts. Overall, this qualitative inspection revealed no observable differences in either morphological appearance (Fig. 5) or cytoskeletal distribution (Suppl. Fig. 1) between real and simulated neurons in any of the cell subclasses, subtypes, and genotypes.

**Figure 5:**
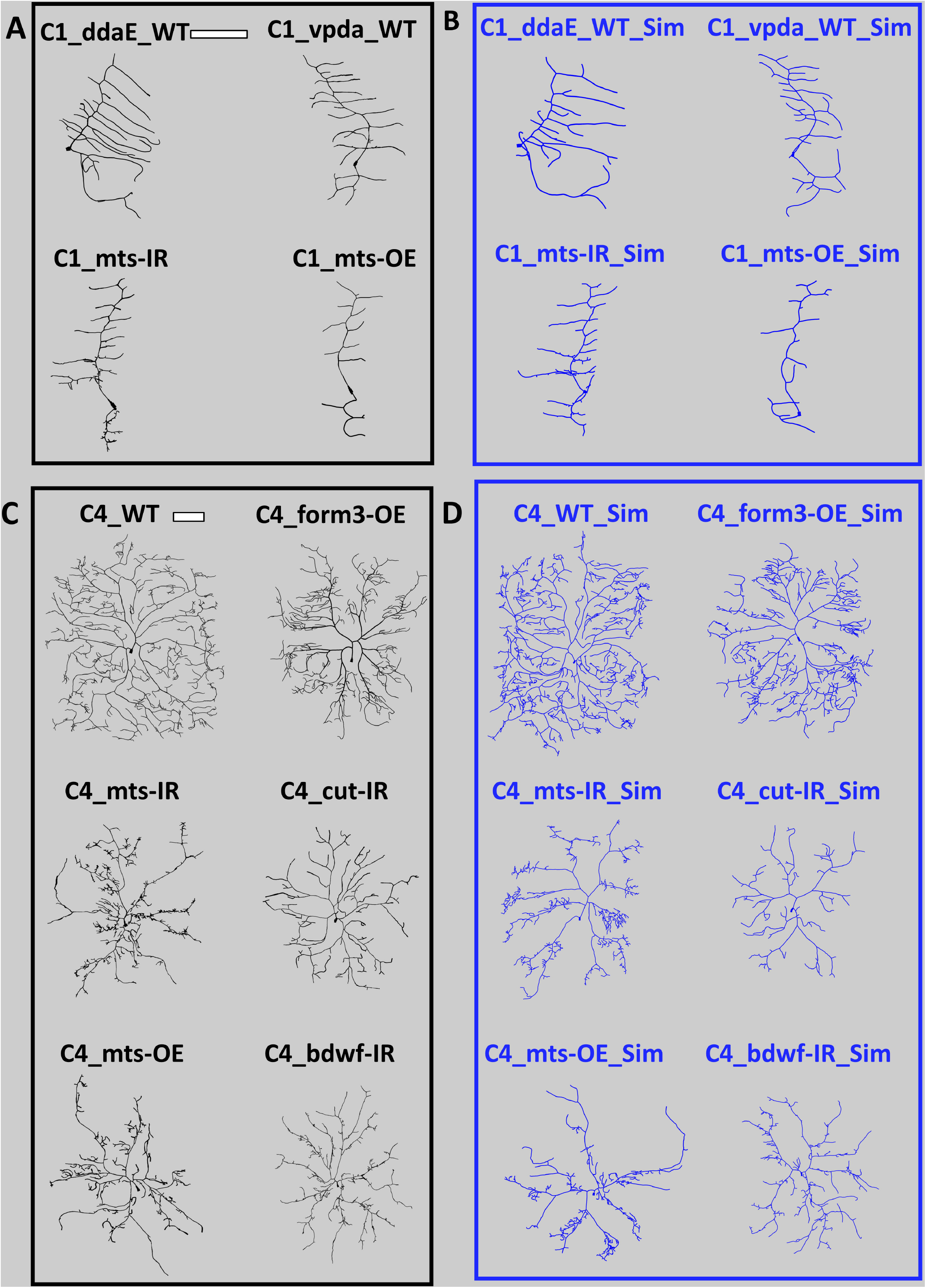
Morphological Comparison of Real and Simulated Neurons. (A) Representative real neurons (black) from the four class I neuron groups (ddaE WT, vpda WT, vpda mts-IR and vpda mts-OE). (B) Representative simulated neurons (blue) from the four class I neuron groups (ddaE WT, vpda WT, vpda mts-IR and vpda mts-OE). (C) Representative real neurons (black) from six Class IV neuron groups (Class IV WT, form3-OE, mts-IR, cut-IR, mts-OE, bdwf-IR). (D) Representative simulated neurons (blue) from six Class IV neuron groups (Class IV WT, form3-OE, mts-IR, cut-IR, mts-OE, bdwf-IR).

We then compared several quantitative morphological parameters between real and simulated neurons from all 17 neuron groups, starting with 10 scalar L-Measure morphometrics properties: number of branches, height, width, total length, maximum path distance, maximum Euclidean distance, maximum branch order, average branch straightness (called “contraction” in L-Measure), average partition asymmetry, and average bifurcation angle. A total of 170 statistical tests demonstrated no significant difference between real and simulated morphologies (Table 3).

**Table 3:** False Discovery Rate-corrected t-test or Wilcoxon test p values of comparisons for 10 L-Measure morphometrics between real and simulated neuron across 17 cell groups.

| Neuron Groups | Number of branch | Height | Width | Length | Max Path Distance | Max Euclidean Distance | Max Branch Order | Contraction | Partition Asymmetry | Local Bifurcation Angle |
| --- | --- | --- | --- | --- | --- | --- | --- | --- | --- | --- |
| C1-ddaE-WT | 1 | 1.000 | 0.994 | 1.000 | 1.000 | 1.000 | 1.000 | 0.993 | 1.000 | 1.000 |
| C1-vpda-WT | 1 | 0.984 | 1.000 | 0.964 | 0.901 | 0.933 | 1.000 | 0.900 | 1.000 | 0.959 |
| C1-vpda-mts-IR | 1 | 1.000 | 1.000 | 1.000 | 0.982 | 0.969 | 1.000 | 1.000 | 1.000 | 0.931 |
| C1-vpda-mts-OE | 0.9367 | 0.981 | 0.770 | 0.883 | 0.918 | 0.785 | 0.791 | 0.946 | 0.683 | 0.953 |
| C4_WT | 1 | 0.970 | 0.850 | 1.000 | 0.940 | 0.762 | 0.788 | 0.734 | 0.820 | 0.910 |
| C4_bdwf-IR | 1 | 0.903 | 0.936 | 0.940 | 0.902 | 0.862 | 0.905 | 0.985 | 0.658 | 1.000 |
| C4_form3-OE | 1 | 0.651 | 0.530 | 0.918 | 0.949 | 0.620 | 0.964 | 1.000 | 0.847 | 0.931 |
| C4_mts-IR | 1 | 0.978 | 0.957 | 0.999 | 0.994 | 0.995 | 0.743 | 0.976 | 0.912 | 0.973 |
| C4_mts-OE | 1 | 0.677 | 0.940 | 1.000 | 0.970 | 0.850 | 0.730 | 1.000 | 1.000 | 0.850 |
| C4_cut_IR | 1 | 0.819 | 0.901 | 0.780 | 0.981 | 0.934 | 0.942 | 0.959 | 0.944 | 0.985 |
| C4_RpL4-IR | 1 | 0.970 | 1.000 | 1.000 | 0.970 | 0.970 | 1.000 | 1.000 | 1.000 | 1.000 |
| C4_RpL17-IR | 1 | 0.660 | 0.846 | 0.920 | 0.974 | 1.000 | 1.000 | 0.937 | 0.909 | 1.000 |
| C4_RpL22-IR | 1 | 1.000 | 1.000 | 1.000 | 1.000 | 1.000 | 1.000 | 0.910 | 0.677 | 0.940 |
| C4_RpL31-IR | 1 | 0.975 | 0.981 | 0.949 | 0.881 | 0.981 | 0.959 | 0.950 | 1.000 | 0.931 |
| C4_RpL35A-IR | 1 | 1.000 | 1.000 | 1.000 | 1.000 | 1.000 | 1.000 | 0.999 | 1.000 | 0.952 |
| C4_RpS10b-IR | 1 | 1.000 | 0.978 | 0.964 | 1.000 | 0.931 | 1.000 | 0.944 | 0.964 | 0.959 |
| C4_RpS24-IR | 0.9592 | 0.965 | 0.834 | 0.924 | 0.644 | 0.969 | 0.578 | 0.930 | 0.784 | 0.805 |

Next, we compared the Sholl-like distribution of total dendritic length against Euclidean distance between the real and the simulated morphologies, and found no observable difference in any of the neuronal groups (Fig. 6). To quantify this comparison, we also employed persistence vectors, a representation of arbor morphology that captures branch distributions similarly to Sholl diagrams but enabling the definition of a formal metric distance (Li et al., 2017). If the real and simulated neurons had different branch distributions, the distance between a real and a simulated neuron would be, on average, larger than the distance between two real or two simulated neurons. The persistence vector arccosine distance was thus measured for each pair of neurons within condition (two real neurons or two simulated neurons) or between condition (one real and one simulated neuron). The within- and between-condition pairwise distances were then compared by t-test, and no significant difference was found for any of 17 neuron groups (Table 4).

**Figure 6:**
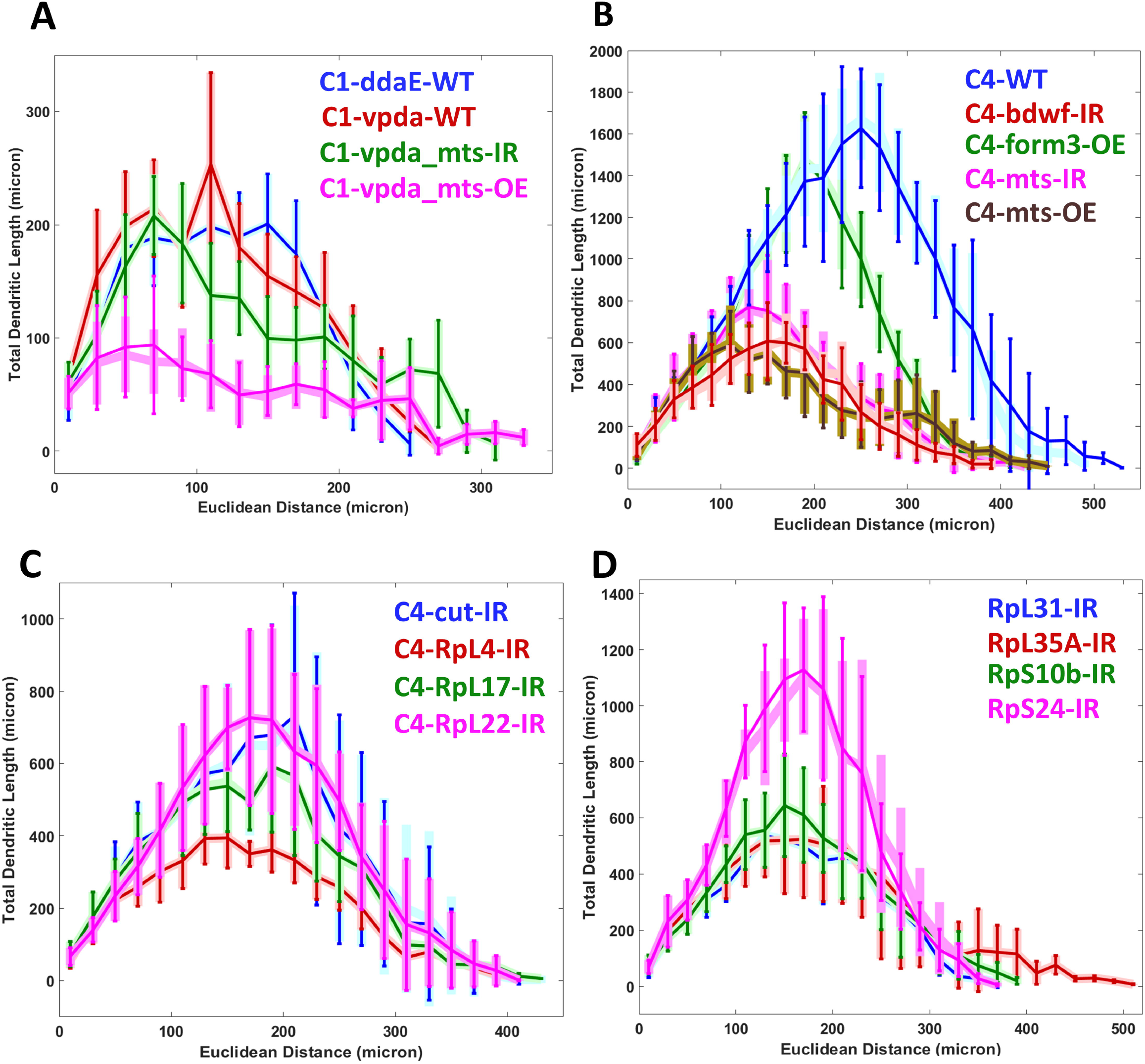
Sholl-Like Comparison Between Real and Simulated Neurons. Sholl-like analysis of total dendritic length against Euclidean distance from the soma for (A) Class I and (B), (C), (D) Class IV neurons. Lighter broader shades of blue, red, green, pink, and brown represent different subclasses and genotypes of the real neurons. Darker thinner lines in analogous colors represent the corresponding subclasses and genotypes of the simulated neurons.

**Table 4:** False Discovery Rate-corrected t-test p values of comparisons for pairwise distances in persistence vectors or global density vectors between within-condition (real vs. real and simulated vs. simulated) and across-conditions (real vs. simulated) for all 17 cell groups. Values greater than 0.1 mean failure to reject the null hypothesis that the real and simulated neurons belong to the same distribution.

| Neuron Group | Persistence Vectos | Arbor Density |
| --- | --- | --- |
| C1-ddaE-WT | 0.9153 | 0.9984 |
| C1-vpda-WT | 0.9301 | 0.989 |
| C1-vpda-mts-IR | 0.9302 | 0.9887 |
| C1-vpda-mts-OE | 0.9134 | 0.9966 |
| C4_WT | 0.9894 | 0.9948 |
| C4_bdwf-IR | 0.9373 | 0.9958 |
| C4_form3-OE | 0.8165 | 0.9938 |
| C4_mts-IR | 0.8821 | 0.999 |
| C4_mts-OE | 0.8442 | 0.998 |
| C4_cut_IR | 0.9075 | 0.9956 |
| C4_RpL4-IR | 0.9941 | 0.9989 |
| C4_RpL17-IR | 0.7161 | 0.9897 |
| C4_RpL22-IR | 0.8507 | 0.9985 |
| C4_RpL31-IR | 0.8196 | 0.9981 |
| C4_RpL35A-IR | 0.8654 | 0.9998 |
| C4_RpS10b-IR | 0.984 | 0.9968 |
| C4_RpS24-IR | 0.9832 | 0.9909 |

Lastly, we comparatively analyzed the overall spatial geometry of the real and simulated morphologies by extracting their arbor density matrices, which capture the dendritic profile of a neuron throughout the invaded region (Cuntz et al., 2010). The average density matrix of each real neuron group was qualitatively similar to that of the corresponding simulated neuron groups in both Class I and Class IV cells (Fig. 7) and for all subtypes and genotypes (Suppl. Fig. 2). To quantify these observations, we vectorized the density matrices of each individual neuron and measured all pairwise arccosine distances, similarly to the persistent vector analysis. Within-condition (real-real or simulated-simulated) distances were not statistically different from between-condition (real-simulated) distances (Table 4).

**Figure 7:**
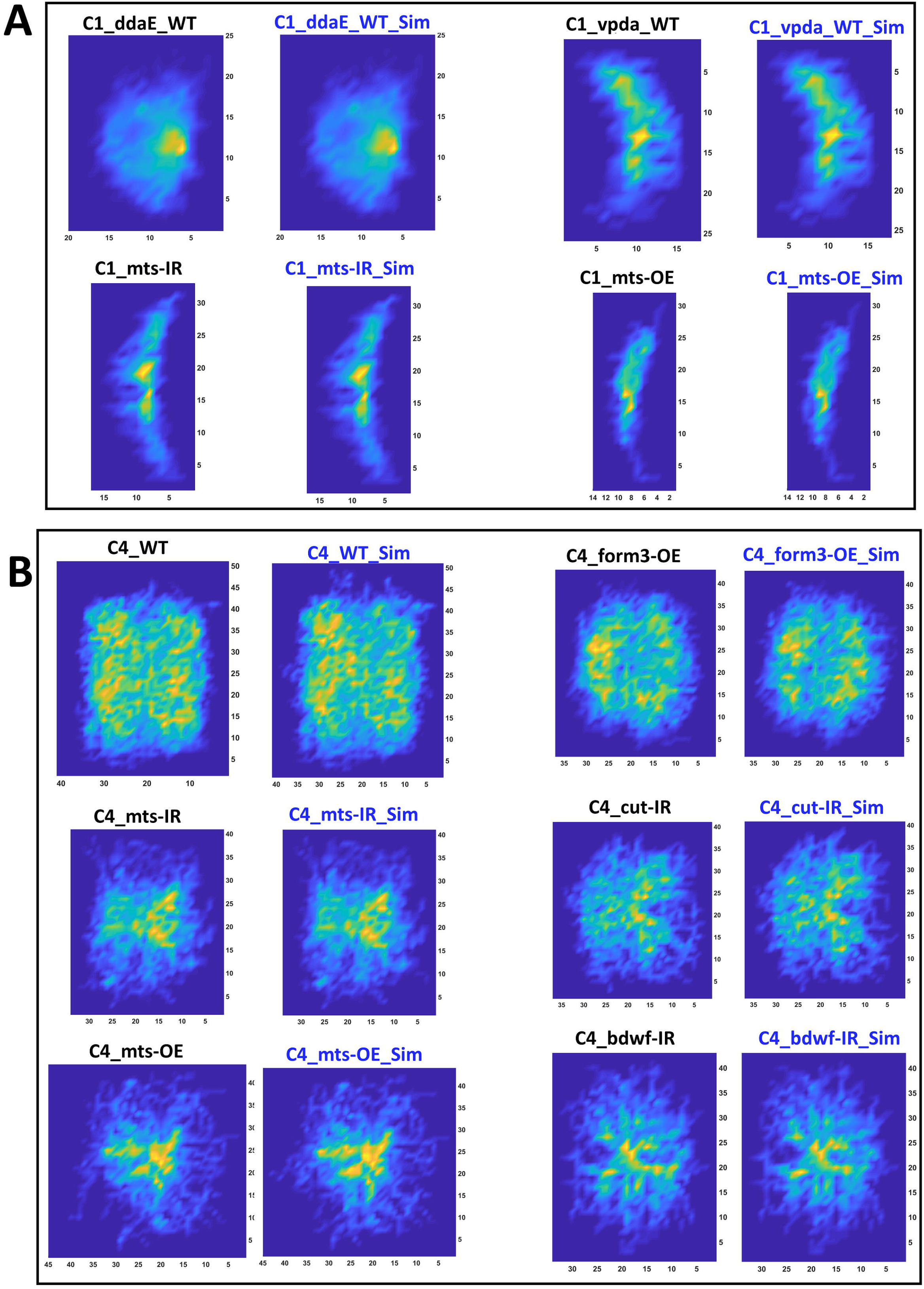
Average Arbor Density of Real and Simulated Neurons. (A) Arbor density averaged across all neurons from each group for Class I neuron types (ddaE WT, vpda WT, vpda mts-IR and vpda mts-OE). Heatmaps for both real (left) and simulated (right) neurons are provided for each group. (B) Arbor density averaged across all neurons from each group for six Class IV neuron groups (WT, form3-OE, mts-IR, cut-IR, mts-OE, bdwf-IR). Heatmaps for both real (left) and simulated (right) neurons are provided for each group. The heatmaps for the 7 remaining Class IV neuron groups are included in Suppl. Fig. 2.

Collectively, these analyses demonstrate the sufficiency of the cytoskeletal determinants in reproducing all geometric features of Class I and Class IV fruit fly sensory da neurons under the control and mutant conditions tested. Class I and Class IV neurons have distinct morphological structures. The mutations in these classes lead to more diverse morphologies and cytoskeletal distribution. However, the influence of microtubule and F-actin is overall consistent through all tested neuron groups: differential distributions of the two cytoskeletal parameters lead to distinct morphological features in the various studied cell classes and genotypes.

## Discussion

*Drosophila* sensory da neurons innervating the barrier epidermis of the larva are a versatile model system to study dendritic morphology. While MT and F-actin have been shown to be directly involved in dendritic growth, maintenance, and plasticity (Conde and Cáceres, 2009; Georges et al., 2008), their precise roles are only recently being elucidated (Nanda et al., 2020). With novel datasets from a diverse set of neuron groups and arbor-wide quantification of MT and F-actin signals, we analyzed the interrelations between cytoskeletal organization and overall dendritic geometry.

Our previous study (Nanda et al., 2020) demonstrated the association between MT and F-actin and the primary causal parameters of dendrogram topology, namely downstream arbor length and branching probability. Specifically, local microtubule quantity predicts the total dendritic length from that location to all the subsequent terminations, and splitting microtubule resources at branch points leads to a division of the arbor length between the corresponding subtrees. At the same time, F-actin enrichment makes a compartment more likely to bifurcate. These two parameters thus fully determine mature arbor size (total dendritic length) and complexity (number of branches). The present study expanded and completed those prior observations by interrelating the local cytoskeleton composition with the spatial geometry of dendritic arbors in addition to their dendrogram topology.

Dendritic orientation is influenced by extracellular guidance cues (Sasaki et al., 2002), and transcriptional mechanisms direct primary dendritic orientation towards active axon terminals and away from inactive axons (Matsui et al., 2013). Prior computational analyses also quantified cell-autonomous relationships between topology and branch angles (Marks and Burke, 2007; Samsonovich and Ascoli, 2003). Here, we identified the influence of cytoskeletal mediator molecules on branch orientation and straightness.

Our results indicate that MT quantity acts as a resistance to dendritic tortuosity and to angular deviation from the previous branches: increased MT levels tended to lead to smaller branch angles across all tested morphological classes, subtypes, and genotypes, as well as to straighter branches in most neuron groups. F-actin was also positively correlated with branch straightness in most neuron groups. However, the correlation of F-actin with branch angle was only significant in a minority of cell subtypes and genotypes and, even among those, the correlation sign switched between positive and negative, suggesting a type-specific influence.

Dendritic arbor generation constrained by MT and F-actin reproduces all geometric properties in both Class I and Class IV neurons across WT da neuron subclasses and a variety of genetic mutations. Notably, the computational model regulates the sampling of topological events, cytoskeletal composition, and spatial orientation solely based on MT and F-actin quantity. Class I and Class IV da neurons lie at the opposite ends of the spectrum of arbor complexity among fruit fly larval da neurons subclasses, and all genetic mutations considered here substantially alter the neuronal arbor relative to their WT counterparts. It is thus remarkable that the same generative model qualitatively and quantitatively captures the dendritic morphology of all tested neuron groups despite their considerable morphological diversity. We interpret this finding by speculating that, among da neurons, distinct morphological classes and genotypes essentially differ in the cytoskeletal composition but share a common developmental program.

Previous studies of basal dendrites in neocortical neurons reported a negative correlation of branch angles with centrifugal branch order (Bielza et al., 2014; Fernandez-Gonzalez et al., 2017). In contrast, we find that branch angles increase with centrifugal branch order in both Class IV and Class I WT neurons. Moreover, in cortical neurons, both dentate gyrus granule cell dendrites (Green and Juraska, 1985) and basal dendrites of pyramidal cells (Bielza et al., 2014) have longer terminal branches than internal branches, whereas we quantified an opposite phenomenon in fly larval da neurons. These contrasting relations could be attributed to the distinct source of input signals on the dendritic arbors of sensory neurons relative to the basal dendrites of cortical neurons, which receive synaptic contacts from local and long-range axons. It would be interesting to examine the cytoskeletal composition of non-sensory neurons to ascertain if it recapitulates the observed differences in the association between branch angles and centrifugal order.

Previous models have used path distance from soma or centrifugal branch order to constrain the simulation of realistic dendrogram topology (Donohue and Ascoli, 2008). The present study demonstrates that cytoskeletal quantity is correlated not only with dendrogram topology, but also with spatial geometry. The computational simulations further corroborate these observations by showing that sampling the experimental cytoskeletal distributions is sufficient to fully reproduce the emergent arbor morphology of real neurons.

Despite this ‘proof by construction’, the model cannot by itself guarantee that the cytoskeletal conditions always cause or even precede the formation of the mature arbor: the cytoskeletal compositions as measured in our data might instead have changed after the arbor reached its final shape. However, experimental evidence from electron microscopy is consistent with lack of MT in terminal branches (Schneider-Mizell et al., 2016). Furthermore, F-actin dynamics have been shown to precede branching events (Nithianandam and Chien, 2018; Sturner et al., 2019) and synaptogenesis (Ritzenthaler et al., 2000). Recent models have also elucidated the role of F-actin dynamics in Class III da neuron branching (Stürner et al., 2022). Pinning down the temporal sequence of cytoskeletal changes underlying dendritic growth will ultimately require time-lapse reconstructions of multi-channel image stacks *in vivo.* That approach involves the meticulous morphological tracing and cytoskeletal quantification from each individual time point, an extremely challenging task at the level of complete arbors (He and Cline, 2011; Rada et al., 2018; Sheintuch et al., 2017).

## Acknowledgment

The authors thank Mahan Mollajafar, Anna Lulushi, Akhil Goel, and Alisha Compton for helping with neuron tracings.

## Author Contribution

GAA and SN designed the study with experimental inputs from DNC. SB provided the experimental images. SN carried out the reconstructions, editing, analysis and simulation. SN drafted the manuscript with help from GAA and all authors edited on the original draft. All authors contributed to editing and finalization. DNC and GAA are responsible for funding acquisition and project management.

## Supplementary Figures

**SF1: Overall morphology along with cytoskeletal distribution of the simulated neurons.** Single representative neurons from 4 Class I neuron types (1 WT subclasses and 2 mutant groups) and 8 Class IV neuron types (1 WT ddaC and 7 mutant groups) are shown, each in two separate images displaying microtubule (red hue) and F-actin (green-blue hue) distributions, respectively. Color bars are constant for all 17 neurons groups. Two separate scale bars for Class I and Class IV neuron groups represent 100 microns.

**SF2: Density profile comparison between the real and simulated neuron groups.** Arbor density averaged across all neurons from each group for the remaining 7 Class IV groups (RpL4-IR, RpL17-IR, RpL22-IR, RpL31-IR, RpL35A-IR, RpS10b-IR and RpS24-IR). Heatmaps for both real (left) and simulated (right) neurons are provided for each group.

## Supplementary Tables

**ST1:** Correlation between branch order and branch angle, average internal branch length, and average terminal branch length for real as well as simulated neurons from all 17 neuron groups.

**ST2:** Correlations between local and remote branch angle, between the angular deviations of sibling branch pairs (D1 and D2), and between maximal angle and branch tilt for all 17 neuron groups.

## References

Akram, M.A., Nanda, S., Maraver, P., Armañanzas, R., Ascoli, G.A., 2018. An open repository for single-cell reconstructions of the brain forest. Sci. Data 5. https://doi.org/10.1038/sdata.2018.6

Bhattacharjee, S., Lottes, E.N., Nanda, S., Golshir, A., Patel, A.A., Ascoli, G.A., Cox, D.N., 2022. PP2A phosphatase regulates cell-type specific cytoskeletal organization to drive dendrite diversity. Front. Mol. Neurosci. 15. https://doi.org/10.3389/FNMOL.2022.926567

Bielza, C., Benavides-Piccione, R., López-Cruz, P., Larrañaga, P., Defelipe, J., 2014. Branching angles of pyramidal cell dendrites follow common geometrical design principles in different cortical areas. Sci. Rep. 4, 1–7. https://doi.org/10.1038/srep05909

Bijari, K., Valera, G., López-Schier, H., Ascoli, G.A., 2021. Quantitative neuronal morphometry by supervised and unsupervised learning. STAR Protoc. 2. https://doi.org/10.1016/J.XPRO.2021.100867

Brown, K.M., Gillette, T.A., Ascoli, G.A., 2008. Quantifying neuronal size: Summing up trees and splitting the branch difference. Semin. Cell Dev. Biol. 19, 485–493. https://doi.org/10.1016/j.semcdb.2008.08.005

Castro, A.F., Baltruschat, L., Stürner, T., Bahrami, A., Jedlicka, P., Tavosanis, G., Cuntz, H., 2020. Achieving functional neuronal dendrite structure through sequential stochastic growth and retraction. Elife 9, 1–38.

Coles, C.H., Bradke, F., 2015. Coordinating neuronal actin-microtubule dynamics. Curr. Biol. 25, R677–91. https://doi.org/10.1016/j.cub.2015.06.020

Conde, C., Cáceres, A., 2009. Microtubule assembly, organization and dynamics in axons and dendrites. Nat. Rev. Neurosci. https://doi.org/10.1038/nrn2631

Corty, M.M., Tam, J., Grueber, W.B., 2016. Dendritic diversification through transcription-factor mediated suppression of alternative morphologies. Development 143, 1351–1362. https://doi.org/10.1242/dev.130906

Cuntz, H., Forstner, F., Borst, A., Häusser, M., 2010. One rule to grow them all: A general theory of neuronal branching and its practical application. PLoS Comput. Biol. 6, e1000877. https://doi.org/10.1371/journal.pcbi.1000877

Das, R., Bhattacharjee, S., Letcher, J.M., Harris, J.M., Nanda, S., Foldi, I., Lottes, E.N., Bobo, H.M., Grantier, B.D., Mihály, J., Ascoli, G.A., Cox, D.N., 2021. Formin 3 directs dendritic architecture via microtubule regulation and is required for somatosensory nociceptive behavior. Dev. 148. https://doi.org/10.1242/DEV.187609/VIDEO-2

Das, R., Bhattacharjee, S., Patel, A.A., Harris, J.M., Bhattacharya, S., Letcher, J.M., Clark, S.G., Nanda, S., Iyer, E.P.R., Ascoli, G.A., Cox, D.N., 2017. Dendritic cytoskeletal architecture is modulated by combinatorial transcriptional regulation in Drosophila melanogaster. Genetics 207, 1401–1421. https://doi.org/10.1534/genetics.117.300393

de la Torre-Ubieta L., Bonni, A., 2011. Transcriptional regulation of neuronal polarity and morphogenesis in the mammalian brain. Neuron 72, 22–40. https://doi.org/10.1016/j.neuron.2011.09.018

Donohue, D.E., Ascoli, G.A., 2008. A Comparative Computer Simulation of Dendritic Morphology. PLoS Comput Biol 4, e1000089. https://doi.org/10.1371/journal.pcbi.1000089

Emoto, K., 2011. Dendrite remodeling in development and disease. Dev. Growth Differ. 53, 277–286. https://doi.org/10.1111/J.1440-169X.2010.01242.X

Falke, E., Nissanov, J., Mitchell, T.W., Bennett, D.A., Trojanowski, J.Q., Arnold, S.E., 2003. Subicular Dendritic Arborization in Alzheimer’s Disease Correlates with Neurofibrillary Tangle Density. Am. J. Pathol. 163, 1615–1621. https://doi.org/10.1016/S0002-9440(10)63518-3

Feng, L., Zhao, T., Kim, J., 2015. neuTube 1.0: A new design for efficient neuron reconstruction software based on the SWC Format. eNeuro 2, ENEURO.0049-14.2014. https://doi.org/10.1523/ENEURO.0049-14.2014

Forrest, M.P., Parnell, E., Penzes, P., 2018. Dendritic structural plasticity and neuropsychiatric disease. Nat. Rev. Neurosci. 2018 194 19, 215–234. https://doi.org/10.1038/nrn.2018.16

Franker, M., Hoogenraad, C., 2013. Microtubule-based transport – basic mechanisms, traffic rules and role in neurological pathogenesis. J Cell Sci 126, 2319–2329. https://doi.org/10.1242/jcs.115030

Georges, P.C., Hadzimichalis, N.M., Sweet, E.S., Firestein, B.L., 2008. The yin-yang of dendrite morphology: Unity of actin and microtubules. Mol. Neurobiol. 38, 270–284. https://doi.org/10.1007/s12035-008-8046-8

Gleeson, P., Davison, A.P., Silver, R.A., Ascoli, G.A., 2017. A Commitment to Open Source in Neuroscience. Neuron 96, 964–965. https://doi.org/10.1016/j.neuron.2017.10.013

Green, E.J., Juraska, J.M., 1985. The dendritic morphology of hippocampal dentate granule cells varies with their position in the granule cell layer: a quantitative Golgi study. Exp. Brain Res. 59, 582–586. https://doi.org/10.1007/BF00261350/METRICS

He, H.Y., Cline, H.T., 2011. Diadem X: Automated 4 Dimensional Analysis of Morphological Data. Neuroinformatics 9, 107–112. https://doi.org/10.1007/s12021-011-9098-x

Hely, T.A., Graham, B., Ooyen, A. V, 2001. A computational model of dendrite elongation and branching based on MAP2 phosphorylation. J. Theor. Biol. 210, 375–84. https://doi.org/10.1006/jtbi.2001.2314

Hu, Y., Roesel, C., Flockhart, I., Perkins, L., Perrimon, N., Mohr, S.E., 2013. UP-TORR: Online tool for accurate and up-to-date annotation of RNAi reagents. Genetics 195, 37–45. https://doi.org/10.1534/GENETICS.113.151340/-/DC1

Iyer, S.C., Wang, D., Iyer, E.P.R., Trunnell, S.A., Meduri, R., Shinwari, R., Sulkowski, M.J., Cox, D.N., 2012. The RhoGEF trio functions in sculpting class specific dendrite morphogenesis in Drosophila sensory neurons. PLoS One 7, e33634. https://doi.org/10.1371/journal.pone.0033634

Jan, Y.-N., Jan, L.Y., 2010. Branching out: mechanisms of dendritic arborization. Nat Rev Neurosci 11, 316–328. https://doi.org/10.1038/nrn2836

Kapitein, L.C., Hoogenraad, C.C., 2015. Building the Neuronal Microtubule Cytoskeleton. Neuron 87, 492–506. https://doi.org/10.1016/j.neuron.2015.05.046

Komendantov, A.O., Ascoli, G.A., 2009. Dendritic excitability and neuronal morphology as determinants of synaptic efficacy. J. Neurophysiol. 101, 1847–1866. https://doi.org/10.1152/JN.01235.2007/ASSET/IMAGES/LARGE/Z9K0040993850012.JPEG

Ledda, F., Paratcha, G., 2017. Mechanisms regulating dendritic arbor patterning. Cell. Mol. Life Sci. 74, 4511–4537. https://doi.org/10.1007/s00018-017-2588-8

Lefebvre, J.L., 2021. Molecular mechanisms that mediate dendrite morphogenesis. Curr. Top. Dev. Biol. 142, 233–282. https://doi.org/10.1016/BS.CTDB.2020.12.008

Lefebvre, J.L., Sanes, J.R., Kay, J.N., 2015. Development of Dendritic Form and Function. Annu. Rev. Cell Dev. Biol. 31, 741–777. https://doi.org/10.1146/annurev-cellbio-100913-013020

Li, Y., Wang, D., Ascoli, G.A., Mitra, P., Wang, Y., 2017. Metrics for comparing neuronal tree shapes based on persistent homology. PLoS One 12, e0182184. https://doi.org/10.1371/JOURNAL.PONE.0182184

Marks, W.B., Burke, R.E., 2007. Simulation of motoneuron morphology in three dimensions. I. Building individual dendritic trees. J. Comp. Neurol. 503, 685–700. https://doi.org/10.1002/cne.21418

Nagel, J.C., Delandre, C., Zhang, Y., Förstner, F., Moore, A.W., Tavosanis, G., 2012. Fascin controls neuronal class-specific dendrite arbor morphology. Development 139, 2999–3009. https://doi.org/10.1242/dev.077800

Nanda, S., Bhattacharjee, S., Cox, D.N., Ascoli, G.A., 2021. An imaging analysis protocol to trace, quantify, and model multi-signal neuron morphology. STAR Protoc. 2, 100567. https://doi.org/10.1016/J.XPRO.2021.100567

Nanda, S., Bhattacharjee, S., Cox, D.N., Ascoli, G.A., 2020. Distinct relations of microtubules and actin filaments with dendritic architecture. iScience 23, 101865. https://doi.org/10.1016/j.isci.2020.101865

Nanda, S., Chen, H., Das, R., Bhattacharjee, S., Cuntz, H., Torben-Nielsen, B., Peng, H., Cox, D.N., Schutter, E. De, Ascoli, G.A., 2018a. Design and implementation of multi-signal and time-varying neural reconstructions. Sci. Data 5, 170207. https://doi.org/10.1038/sdata.2017.207

Nanda, S., Das, R., Bhattacharjee, S., Cox, D.N., Ascoli, G.A., 2018b. Morphological determinants of dendritic arborization neurons in Drosophila larva. Brain Struct. Funct. 223, 1107–1120. https://doi.org/10.1007/s00429-017-1541-9

Nanda, S., Das, R., Cox, D.N., Ascoli, G.A., 2017. Structural Plasticity in Dendrites: Developmental Neurogenetics, Morphological Reconstructions, and Computational Modeling, in: Laura Petrosini (Ed.), Neurobiological and Psychological Aspects of Brain Recovery. Springer Press, Contemporary 753 Clinical Neuroscience Series: Cham. 30, pp. 1–34. https://doi.org/10.1007/978-3-319-52067-4_1

Nithianandam, V., Chien, C.T., 2018. Actin blobs prefigure dendrite branching sites. J. Cell Biol. 217, 3731–3746. https://doi.org/10.1083/jcb.201711136

Parekh, R., Ascoli, G.A., 2015. Quantitative investigations of axonal and dendritic arbors: development, structure, function, and pathology. The Neuroscientist: a review. J. bringing Neurobiol. Neurol. psychiatry 21, 241–254. https://doi.org/10.1177/1073858414540216

Rada, L., Kilic, B., Erdil, E., Ramiro-Cortés, Y., Israely, I., Unay, D., Cetin, M., Argunsah, A.Ö., 2018. Tracking-assisted Detection of Dendritic Spines in Time-Lapse Microscopic Images. Neuroscience 394, 189–205. https://doi.org/10.1016/j.neuroscience.2018.10.022

Ritzenthaler, S., Suzuki, E., Chiba, A., 2000. Postsynaptic filopodia in muscle cells interact with innervating motoneuron axons. Nat. Neurosci. 3, 1012–1017. https://doi.org/10.1038/79833

Ropireddy, D., Ascoli, G.A., 2011. Potential synaptic connectivity of different neurons onto pyramidal cells in a 3D reconstruction of the rat hippocampus. Front. Neuroinform. 5, 5. https://doi.org/10.3389/FNINF.2011.00005/BIBTEX

Samsonovich, A. V., Ascoli, G.A., 2005. Statistical determinants of dendritic morphology in hippocampal pyramidal neurons: A hidden Markov model. Hippocampus 15, 166–183. https://doi.org/10.1002/hipo.20041

Samsonovich, A. V., Ascoli, G.A., 2003. Statistical morphological analysis of hippocampal principal neurons indicates cell-specific repulsion of dendrites from their own cell. J. Neurosci. Res. 71, 173–187. https://doi.org/10.1002/jnr.10475

Schindelin, J., Arganda-Carreras, I., Frise, E., Kaynig, V., Longair, M., Pietzsch, S., Rueden, C., Saalfeld, S., Schmid, B., Tinevez, J.Y., Pietzsch, T., Preibisch, S., Rueden, C., Saalfeld, S., Schmid, B., Tinevez, J.Y., White, D.J., Hartenstein, V., Eliceiri, K., Tomancak, P., Cardona, A., 2012. Fiji: an open-source platform for biological-image analysis. Nat Methods 9, 676–682. https://doi.org/10.1038/nmeth.2019

Schneider-Mizell, C.M., Gerhard, S., Longair, M., Kazimiers, T., Li, F., Zwart, M.F., Champion, A., Midgley, F.M., Fetter, R.D., Saalfeld, S., Cardona, A., 2016. Quantitative neuroanatomy for connectomics in Drosophila. Elife 5, e12059. https://doi.org/10.7554/eLife.12059

Scorcioni, R., Polavaram, S., Ascoli, G.A., 2008. L-Measure: a web-accessible tool for the analysis, comparison and search of digital reconstructions of neuronal morphologies. Nat. Protoc. 3, 866–876. https://doi.org/10.1038/nprot.2008.51

Sheintuch, L., Rubin, A., Brande-Eilat, N., Geva, N., Sadeh, N., Pinchasof, O., Ziv, Y., 2017. Tracking the Same Neurons across Multiple Days in Ca2+ Imaging Data. Cell Rep. 21, 1102–1115. https://doi.org/10.1016/j.celrep.2017.10.013

Stiso, J., Bassett, D.S., 2018. Spatial Embedding Imposes Constraints on Neuronal Network Architectures. Trends Cogn. Sci. 22, 1127–1142. https://doi.org/10.1016/J.TICS.2018.09.007

Stürner, T., Ferreira Castro, A., Philipps, M., Cuntz, H., Tavosanis, G., 2022. The branching code: A model of actin-driven dendrite arborization. Cell Rep. 39, 110746. https://doi.org/10.1016/j.celrep.2022.110746

Sturner, T., Tatarnikova, A., Mueller, J., Schaffran, B., Cuntz, H., Zhang, Y., Nemethova, M., Bogdan, S., Small, V., Tavosanis, G., 2019. Transient localization of the arp2/3 complex initiates neuronal dendrite branching in vivo. Dev. 146, dev171397. https://doi.org/10.1242/dev.171397

