## Supplementary figures and images for "Local microtubule and F-actin distributions fully determine the spatial geometry of *Drosophila* sensory dendritic arbors"

### Suppl. Fig. 1

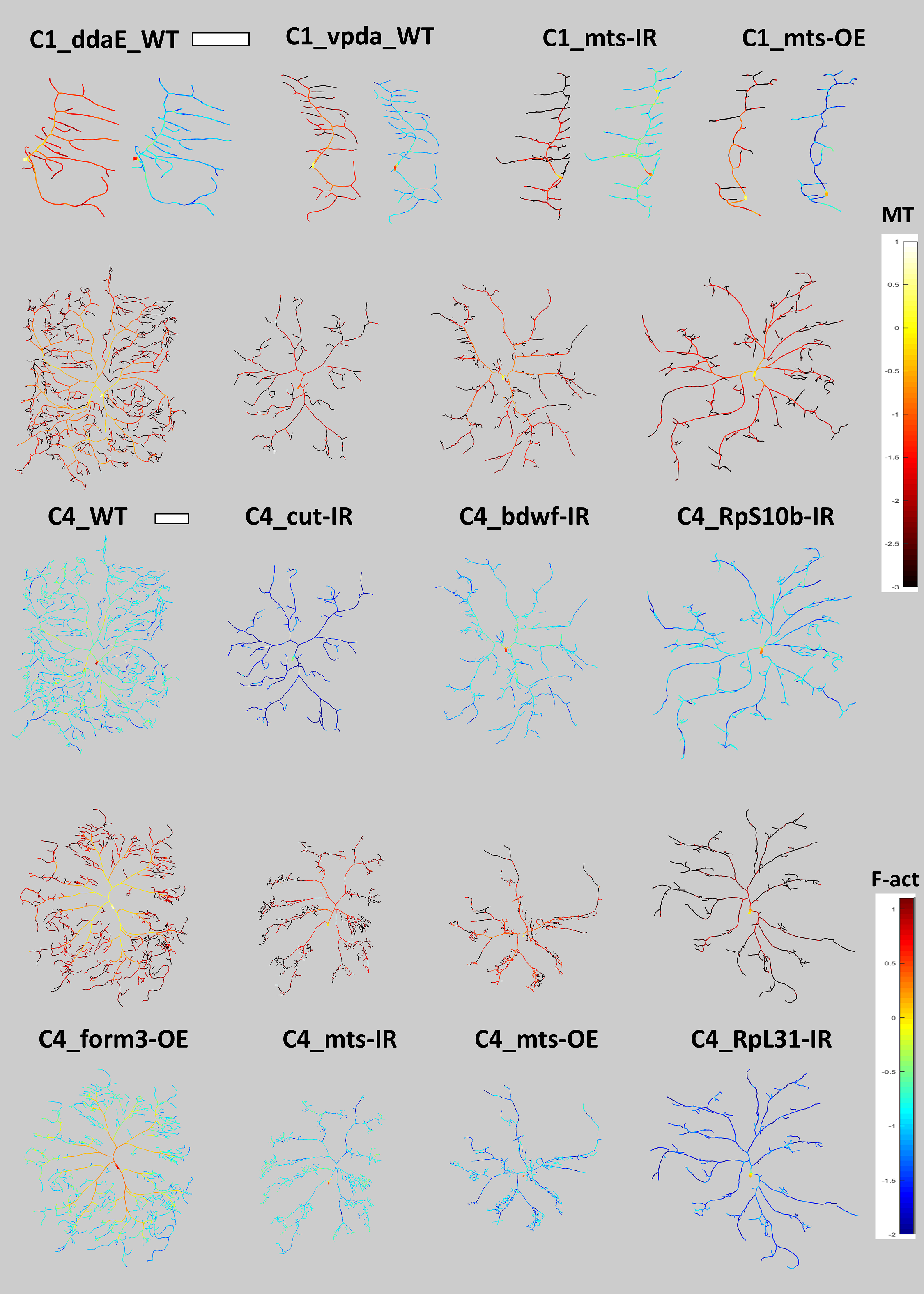

### Suppl. Fig. 2

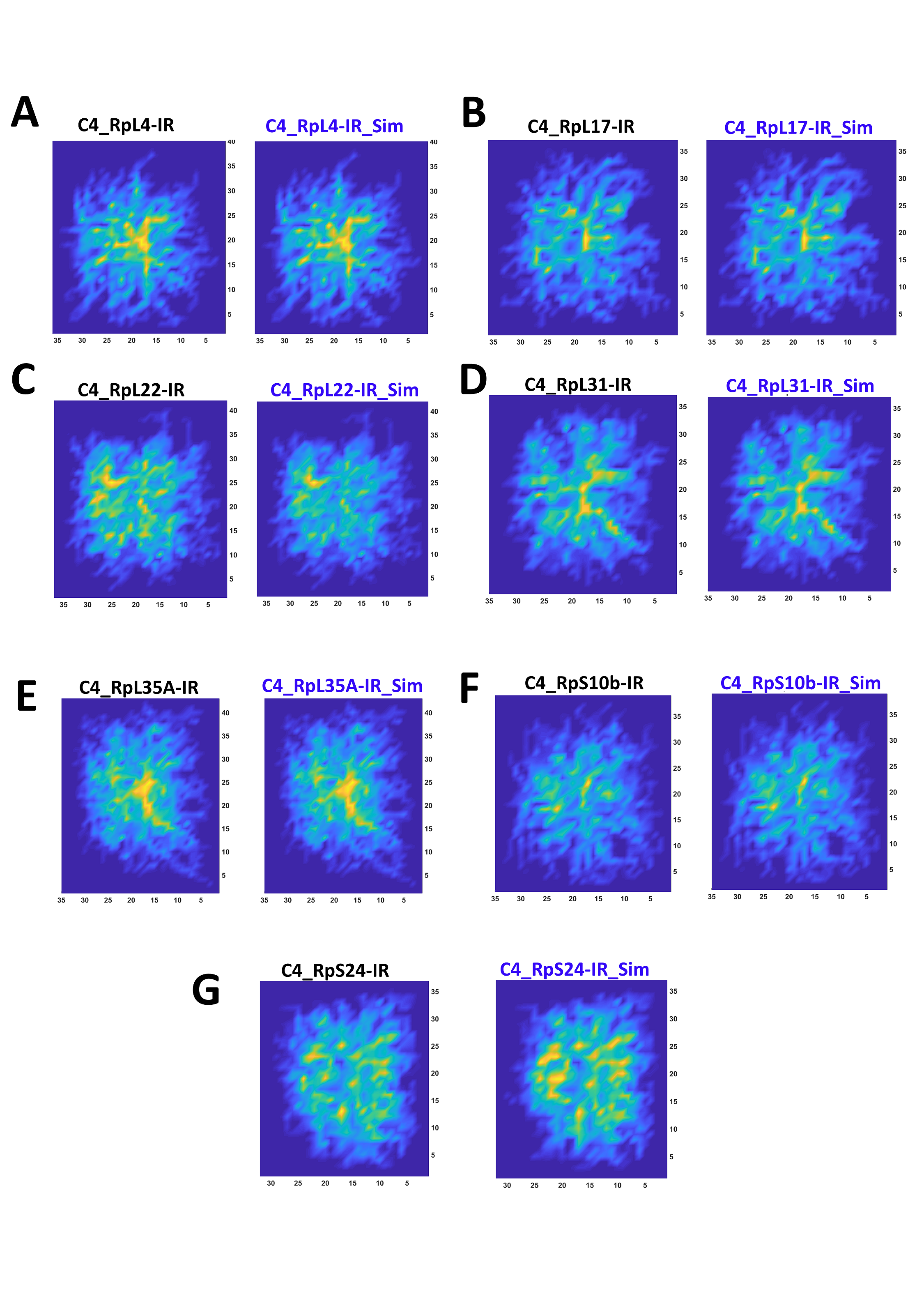
